# Molecular-Dynamics Simulation Methods for Macromolecular Crystallography

**DOI:** 10.1101/2022.04.04.486986

**Authors:** David C. Wych, Phillip C. Aoto, Lily Vu, Alexander M. Wolff, David L. Mobley, James S. Fraser, Susan S. Taylor, Michael E. Wall

**Affiliations:** Computer, Computational, and Statistical Sciences Division, Los Alamos National Laboratory, Los Alamos, NM, 87545, USA; Center for Nonlinear Studies, Los Alamos National Laboratory, Los Alamos, NM, 87545, USA; Department of Pharmaceutical Sciences, University of California, Irvine, CA, 92697, USA; Department of Pharmacology, University of California, San Diego, La Jolla, CA, 92093; Department of Bioengineering and Therapeutic Sciences, University of California, San Francisco, CA, 94158, USA; Department of Chemistry, University of California, Irvine, CA, 92697, USA; Department of Chemistry and Biochemistry, University of California, San Diego, La Jolla, CA, 92093

**Keywords:** Molecular-dynamics simulations, macromolecular crystallography, electron density, water structure, protein structure, conformational ensemble, protein kinase

## Abstract

To assess the potential benefits of molecular-dynamics (MD) simulations for macromolecular crystallography (MX), we performed room-temperature X-ray diffraction studies of the catalytic subunit of mouse protein kinase A (PKA-C). We then performed crystalline MD simulations of PKA-C, computed simulated electron densities from the water, protein, and ion components of the MD simulations, and carefully compared them to the initial crystal structure. The results led to the development of an MD-MX analysis procedure and several associated methods: 1) *density comparison* to evaluate consistency between the MD and the initial crystal structure model; 2) *water building* to generate alternative solvent models; and 3) *protein remodeling* to improve the crystal structure where interpretation of density is unclear. This procedure produced a revised structure of PKA with a new ordered water model and a modified protein structure. The revisions yield new insights into PKA mechanisms, including: a sensitivity of the His294 conformation to protonation state, with potential consequences for regulation of substrate binding; a remodeling of the Lys217 side chain along with a bound phosphate; an alternative conformation for Lys213 associated with binding to the regulatory subunit; and an alternative conformation for catalytic base Asp166 and nearby waters, suggesting a mechanism of progression of the phosphotransfer reaction via changes in Mg^2+^ coordination. Based on the benefits seen applying these methods to PKA, we recommend incorporating our MD-MX procedure into MX studies, to decide among ambiguous interpretations of electron density that occur, inevitably, as part of standard model refinement.

## Introduction

Macromolecular crystallography (MX) has produced most of what is known about the atomic structure of proteins (1). Historically, protein crystal structures have consisted of a single set of atomic coordinates (mean positions), temperature factors (positional variance), and occupancies (fraction of the crystal where the atom is present). Structural models with just a single set of parameters for each atom, however, are limited in their ability to describe the full range of conformational variations that may be present in protein crystals. A number of approaches have been developed over the years to overcome this limitation, e.g. allowing for crystal structure models with multiple conformations of selected side chains (2, 3), or ensemble models with multiple copies of the entire protein (4–6). The resulting improvements in descriptions of conformational heterogeneity are needed to understand biological mechanisms involved in catalysis, molecular recognition, and allostery (7).

Although multi-conformer modeling has improved the ability of protein crystallography to describe conformational heterogeneity, methods can fail in regions where crystallographic density is difficult to interpret, as can occur, e.g., at the protein-solvent interface. At the interface, protein atoms might be built into a region that, in the actual crystal, is mainly occupied by solvent. Similarly, standard methods for building ordered waters into a crystal structure (e.g. water picking (8)) make use of a supplied protein model; if this model contains errors, these methods can place waters into a region mainly occupied by protein. Addressing these issues might increase crystal structure accuracy; in particular, improving the water structure has been highlighted as a route to improved model accuracy (9). Improved water structure models also can improve the modeling of molecular recognition and ligand binding (10, 11), and yield insights into the regulation of protein dynamics and allostery (12) and the formation of protein-protein binding interfaces (13). The hydration layer at the surface of proteins has been shown to be central to an understanding of protein structure, dynamics, and function (14).

Because crystallographic density does not come with labels, independent information is sometimes required to help distinguish the protein and solvent components. Recent developments suggest that molecular dynamics (MD) simulations might be a promising means of obtaining such information. MD is a powerful computational method for providing insight into biomolecular structure and dynamics. In MD, a set of coordinates and potential energy parameters are used to model the atoms in a molecular system. The dynamics of the system are computed using Newtonian numerical integration, resulting in a trajectory that can be analyzed to capture phenomena on a time scale and at a resolution that is often inaccessible by laboratory measurement. Techniques borrowed from MD have been used for many years to sample conformations in crystallographic refinement (6, 8). In addition, crystalline MD simulation methods have advanced substantially in recent years, in large part thanks to MD studies of diffuse X-ray scattering (15–24). Recent studies have shown that crystalline MD simulations can reproduce both the positions of ordered waters (25) and the B-factors from crystallographic refinement (22, 26). These advances suggest that crystalline MD techniques have developed to the point where they might benefit macromolecular crystallography workflows.

Here we investigate the utility of MD simulations in interpreting crystallographic density. Our MD approach is grounded in methods developed in a study that investigated the ability of MD simulations to recover crystallographic water structure (25). In that study, it was found that crystalline MD simulations of endoglucanase were able to reproduce the positions of nearly all the ordered waters from combined neutron- and X-ray-crystallography experiments, but only when protein heavy atoms were harmonically restrained to the crystal structure. The restraints biased the protein atoms toward the coordinates from single-structure experimental refinement and suppressed dynamics associated with anharmonic motions and structural heterogeneity. This left open the question of whether applying restraints to an *ensemble*, rather than a single structure, might improve the simultaneous modeling of protein conformational heterogeneity and the water network. To address this question, we have developed an MD-MX procedure for using ensemble-restrained MD simulations to revise the protein and water model in MX structures.

The system we studied in developing this procedure is the catalytic subunit of mouse protein kinase A (PKA), a cyclic adenosine monophosphate (cAMP) dependent protein kinase involved in the regulation of fundamental biological processes that include metabolism, development, memory, and immune response. PKA exists in the cell as a tetramer consisting of two heterodimers with each dimer containing a catalytic (PKA-C) and regulatory (PKA-R) subunit. PKA-C (kinase domain) is activated by phosphorylation on its activation loop by PDK1 or via autophosphorylation as it comes off of the ribosome (27). In addition, because intracellular concentrations of ATP are high (mM) relative to ADP (~10^3^-fold) and PKA (~10^6^-fold), in cells the active ATP-bound form of PKA-C would be highly favored, and the enzyme would be constitutively signaling. To prevent this, PKA-R binds to the active site groove of PKA-C, blocking activity and phosphorylation of partner proteins, and making PKA activity dependent on the second messenger, cAMP. Binding of cAMP to PKA-R releases it from PKA-C, unleashing the catalytic activity. In our system, a small (20 amino acid) pseudo-substrate peptide (IP20) from protein kinase inhibitor (PKI) blocks the active site. PKI competes with PKA-R binding to the inhibitor site in certain tissues and leads to nuclear export (28).

PKA-C has been extensively studied using crystallography, neutron diffraction (29), nuclear magnetic resonance (30), and other methods, due to its biological significance and role as a prototypical kinase (31). There is also evidence of biologically important dynamics and allostery making it a prime choice for study using crystalline MD (30). Cryo-cooling crystals, though a common practice to reduce radiation damage, suppresses dynamics and traps proteins in conformations that may not be representative of their true structural ensembles and interaction networks *in vivo* (32). To capture PKA-C dynamics, therefore, crystals were grown at room temperature and X-ray crystallographic diffraction data were collected at 12°C. Crystal structures were obtained using both single-structure and ensemble refinement. Most likely due to the room temperature crystallization conditions, we found that adenosine triphosphate (ATP), which was used during the purification and crystallization and is often present in the active site in other PKA-C crystal structures, had hydrolyzed, allowing us to capture the immediate products of hydrolysis --- adenosine diphosphate (ADP) and inorganic phosphate. Such hydrolysis had not been observed previously when crystals were grown at 4°C and data collected at cryogenic temperatures.

Our study of PKA-C structure and function guided the development of the MD-MX procedure. The procedure involves three methods: (1) *density comparison*, enabling direct comparison with experimental data for assessment of the consequences of alternative modeling choices, including protonation states; (2) *water building*, which calculates separate structure factors for the protein and water components of the simulation and produces an alternative ordered water model to the one generated by crystallographic refinement; and (3) *protein remodeling*, which uses both the MD density and trajectory snapshots to improve the modeling of residues where interpretation of the density is unclear. Combining all these methods yielded a revised structural model with implications for PKA biology. In producing this model, we found a sensitivity of the conformation of His294 to protonation state, yielding insight into the role of this residue in the modulation of substrate binding affinity (33). The revised model includes a different conformation for Lys217 and introduces a coordinated free phosphate nearby. It also includes multi-conformer states for: (i) Lys213, including a conformation associated with binding to the regulatory domain; and (ii) the active site catalytic base Asp166, including a conformation that appears to be associated with the progression of the phosphotransfer reaction. Based on the benefits seen applying these methods to PKA-C, we recommend incorporating our MD-MX procedure into MX studies, to produce multi-conformer models and to decide among ambiguous interpretations of electron density that occur, inevitably, as part of standard model refinement. The resulting models may yield biological insights beyond what standard crystallographic modeling techniques currently provide.

## Results

### MD-MX analysis procedure

Our main goal in this study was to determine whether crystalline MD simulations can provide insights into protein structure beyond what is available using standard protein crystallography tools. To this end, we developed an MD-MX procedure for using crystalline MD simulations to revise crystal structures (Fig. 1). A distinguishing feature of this procedure is the calculation of structure factors and electron densities from MD trajectories. Such calculations previously enabled quantitative comparisons to assess the agreement of MD simulations with dynamics (21–23) and ordered waters (25) in protein crystals. Here we extend these capabilities to improve interpretation of crystallographic density.

**Figure 1:**
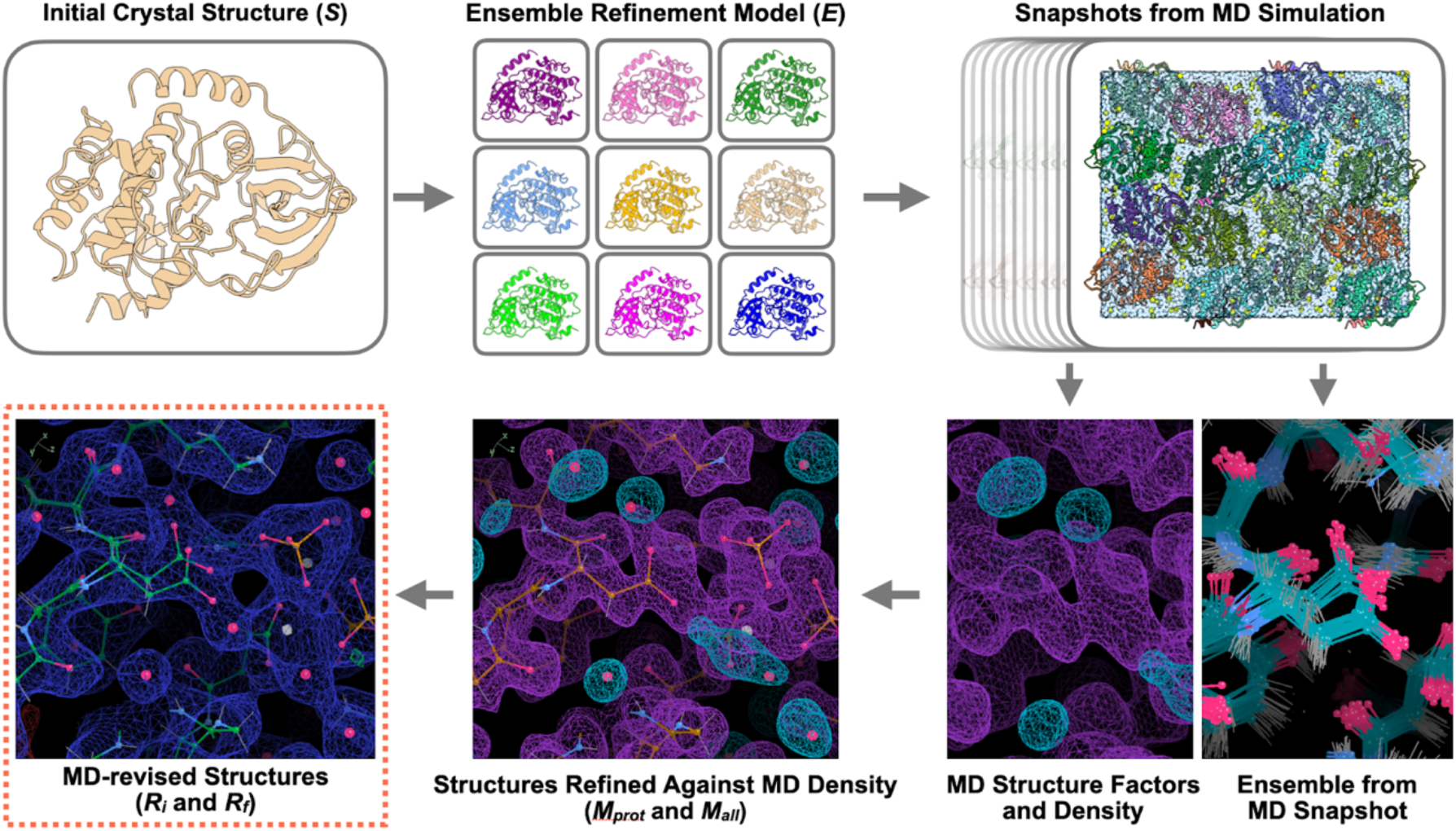
MD-MX data analysis procedure. In this procedure, an initial crystal structure, *S* (top left) is used to prepare a MD simulation model, using ensemble refinement to generate a set of diverse conformations (the Ensemble Refinement Model, *E*; top center). Snapshots from the MD simulation (top right) are used to generate simulated data and MD ensemble visualizations (bottom right). The initial structure is refined against the simulated data, leading to a revised protein structure, *M_prot_*, and a new water network, *M_all_* (bottom center). The revised structure is refined against the experimental data, to produce the initial MD-revised structure *R_i_*, and manual improvements are made using insights from the MD density and ensemble, leading to a final MD-revised structure, *R_f_* (bottom left).

The MD-MX procedure was applied to crystalline PKA-C using X-ray data collected at a temperature of 12°C (Methods); the key steps are illustrated in Figure 1. First, an initial crystal structure (*S*) was obtained using molecular replacement and refinement adding ordered waters (R-factors in Tables 1, S1). To generate an MD model with realistic conformational variation, the structure was further subjected to ensemble refinement, producing an ensemble refinement model (*E*) with 32 protein structures (R-factors in Tables 1, S1). These structures were randomly packed to assemble a crystalline system of 2×2×2 unit cells with four proteins per unit cell, and the resulting system was prepared for restrained MD simulations, in which the heavy atom positions were biased toward the positions in *E* (Methods). After initial equilibration, the protein structure differed from the crystal structure; to provide sufficient time for the protein to relax into a configuration that was close to the crystal structure, we used the final 10 ns of the full 100 ns trajectory for analysis. Snapshots were recorded every 2 ps and were used to produce ensemble visualizations and mean structure factors and densities from protein, solvent, and ion components. The initial structure *S* was refined first against the non-solvent MD structure factors, and then against the total MD structure factors, producing structures *M_prot_* and *M_all_*, respectively. The *M_all_* structure was refined against experimental data, producing an initial MD-revised structure *R_i_*. Lastly, information from the density and MD ensembles were used to remodel the structure by hand, yielding the final MD-revised structure *R_f_*.

**Table 1:**
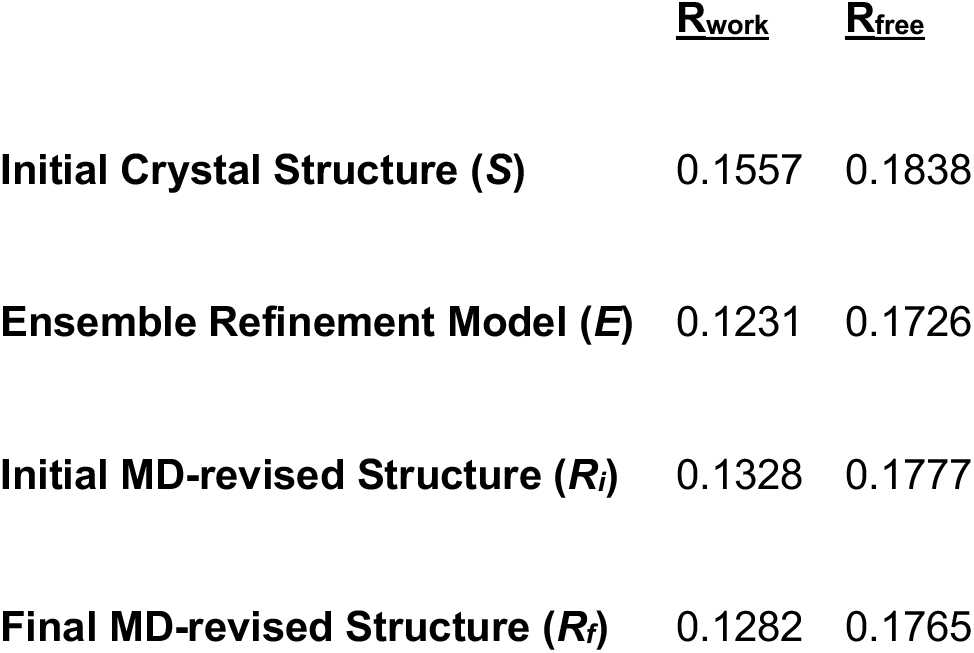
R-factors for structures refined against experimental X-ray data.

Applying the MD-MX procedure to PKA-C produced new information and insights, presented here. The presentation is organized into sections, according to the method that was used to obtain the information. The following sections present the methods, along with the key results they produced.

### Density comparison to assess agreement between MD and crystal structures

The overlap between the MD density and the initial structure *S* is satisfactory for most of the protein. As an exception, two histidines (His62 and His294) showed changes in the density depending on the protonation states. The side chain density for both agreed with *S* when singly protonated on the epsilon nitrogen, as in a neutron crystal structure (PDB ID: 6e21 (29)) (Fig. 3A). When doubly protonated, however, the His62 imidazole ring rotates 45 degrees, and the His294 side chain enters a new rotameric state, which makes room for two ordered waters in the neighborhood (Fig. 2A and 2C).

**Figure 2:**
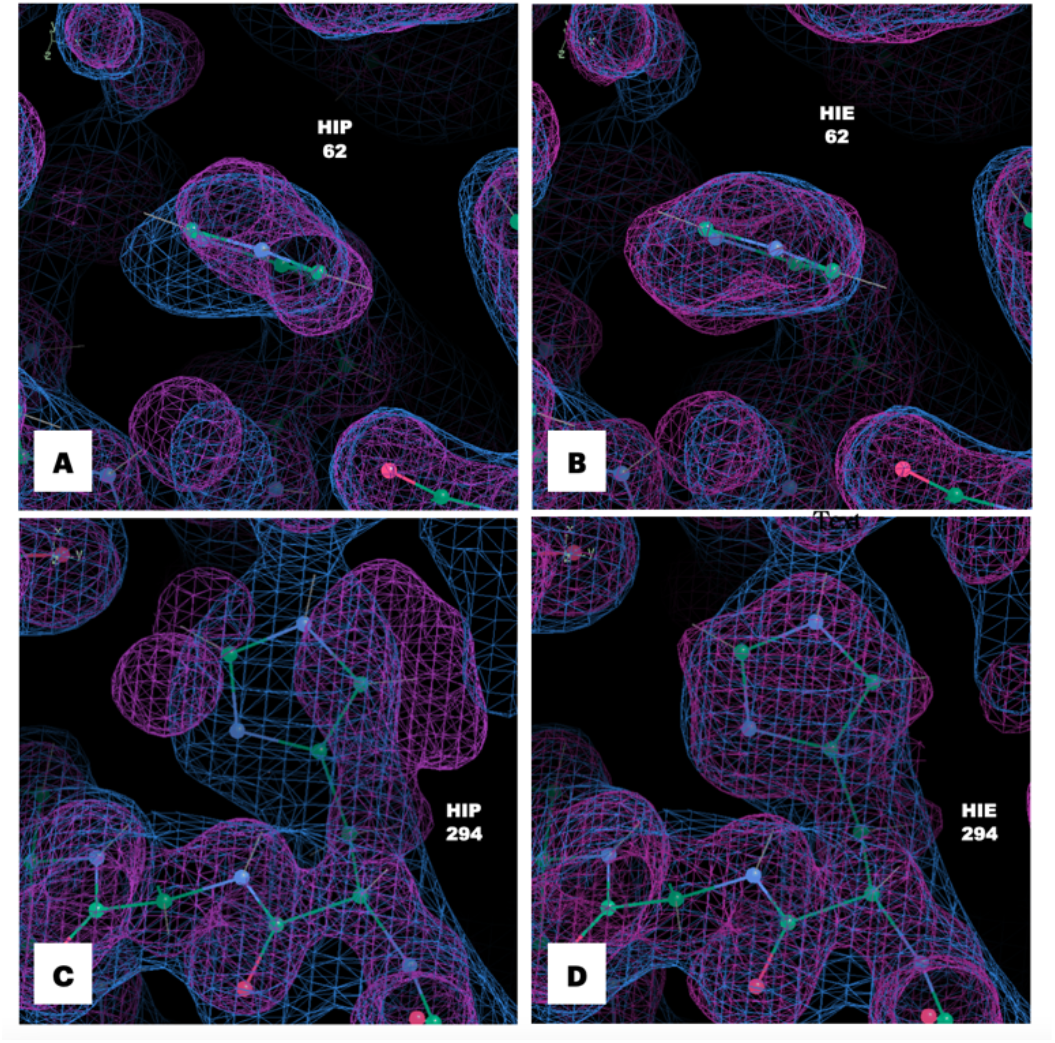
MD densities for His62 and His294 are sensitive to protonation state. The densities agree with the crystallographic density when the residues are epsilon protonated, but disagree when they are doubly protonated. Coordinates of *S* are shown as sticks, 2F_o_-F_c_ in blue (1 sigma isosurface), and total density from 90-100ns segment of a 200 kJ mol^-1^ nm^-2^ MD simulation in pink (1 sigma isosurface). **A:** Histidine 62: MD density from doubly-protonated (HIP) simulation. **B:** Histidine 62: MD density from simulating with histidine singly-protonated (on epsilon nitrogen; HIE) **C:** Histidine 294: MD density from doubly-protonated (HIP) simulation. **D:** Histidine 294: MD density from simulating with histidine singly-protonated (on epsilon nitrogen; HIE).

**Figure 3:**
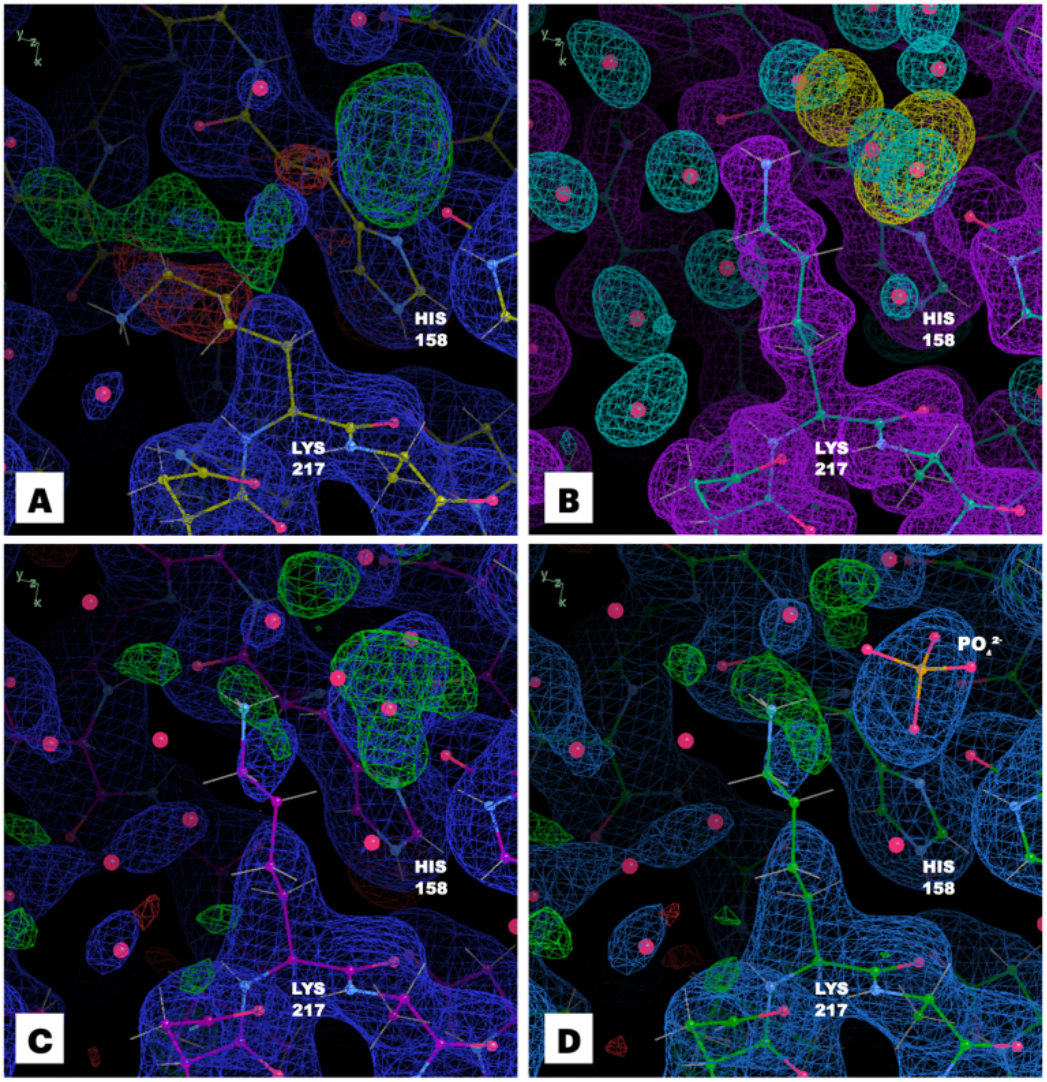
MD remodeling improves interpretation of density in the neighborhood of lysine 217. **A:** coordinates, 2F_o_-F_c_ (blue, 1 sigma isosurface) and F_o_-F_c_ (positive in green and negative in red, 3 sigma isosurface) from *S*. **B:** coordinates from *M*_all_, with MD protein (purple, 1 sigma isosurface), solvent (blue, 3 sigma isosurface), and chloride density (yellow, 10 sigma isosurface) from the 90-100ns segment of the simulation; the MD simulation suggests a different conformation for the side chain and a number of ordered waters; it also includes a spot of chloride density in the same position as positive difference density in panel A. **C:** coordinates, 2F_o_-F_c_ (blue, 1 sigma isosurface) and F_o_-F_c_ (positive in green and negative in red, 3 sigma isosurface) from *R*_i_; the shape of the difference density is suggestive of a coordinated free phosphate molecule. **D:** coordinates, 2F_o_-F_c_ (blue, 1 sigma isosurface) and F_o_-F_c_ (positive in green and negative in red, 3 sigma isosurface) from model *R_f_*; the revised side chain conformation, water network, and phosphate are plausible and improve the difference density in the region. His158 is also shown as a reference point.

To assess the agreement between the MD and crystallographic waters, MD densities were computed using just the water atoms. Peak-finding produced a picture of possible positions of ordered waters; to limit the number of peaks to those more likely to be significant, peaks where the density exceeds 1 electron per cubic angstrom were selected for comparisons with waters from the initial crystal structure (Methods). Recovery of crystallographic waters was assessed using a recall statistic (fraction of crystallographic waters that have a MD peak nearby): the simulation reproduces the positions of 137 of 148 (92.6%) of the crystallographic waters to within 1 Å, similar to a prior MD study of endoglucanase (25).

### Water building to generate an alternative solvent model

We used the water density to develop an alternative ordered water model for the crystal structure. To aid in the disambiguation of protein and solvent density, we performed “protein-first refinement”. The initial crystal structure *S*, stripped of its waters, was refined against the structure factors computed only from the non-solvent components of the simulation, producing *M*_prot_. Next, the *M*_prot_ model was used as an input for refinement against the structure factors computed from all atoms in the simulation, including the waters (Methods). In this second refinement, waters were added using Phenix. The resulting model, *M*_all_, was used as an initial structure for refinement against experimental data. In this refinement, the waters in *M*_all_ were preserved, and no new waters were added. The resulting initial MD-revised structure, *R_i_*, thus contains waters purely derived from the MD simulation. The R-work and R-free values for *R_i_* (0.1328, 0.1777) decreased substantially compared to *S* (0.1557, 0.1838) (Table 1, rows 1 and 3 and Table S1). The *M*_all_ structure had 494 waters (compared with 148 from *S*). The ordered water model in *M*_all_ reproduced 136 of 148 (91.9%) of the waters in *S* to within 1 Å, which is very similar to the value observed using peak-finding above (137 of 148).

### Protein remodeling where interpretation of density is unclear

The water building method yielded a revised crystal structure *R_i_* with an alternate water model and improved R-factors compared to the initial crystal structure *S*. The *R_i_* structure, however, still had regions that needed improvement. To address these issues, we developed the protein remodeling method.

Protein remodeling uses MD density and ensemble snapshots to guide further revisions to the structure resulting from the water building step. The MD density is used to determine where there might be an alternate conformation or a different way to build a side chain, and to suggest where small molecules might be built into the model. The MD ensemble snapshots suggest specific conformations for rebuilding the side chain, as starting points for further rounds of crystallographic refinement. In applying these steps to the *R_i_* structure, the revisions slightly improved the agreement with the data: the final MD-revised structure *R_f_* had an R_work_ of 0.1282 and an R_free_ of 0.1765, compared to values of 0.1328 and 0.1777, respectively, for the *R_i_* structure (Table 1). Importantly, the resulting structure *R_f_* yielded new insights into mechanisms of PKA activity and regulation (Discussion).

We first describe the revisions in the modeling of Lys217. In structure *S*, Lys217 shows negative difference density at the end of the side chain, and positive difference density extending from the beta carbon (Fig. 3A). In the ensemble structure *E*, side chain atoms have diverse conformations, some of which are close to that in structure *S*, but most of which are very different (Fig. 4A). Many of the individual structures in *E* have the side chain positioned so that the amino group falls within a spot of density and that is far from the side chain atoms in *S* (Fig. 4A).

**Figure 4:**
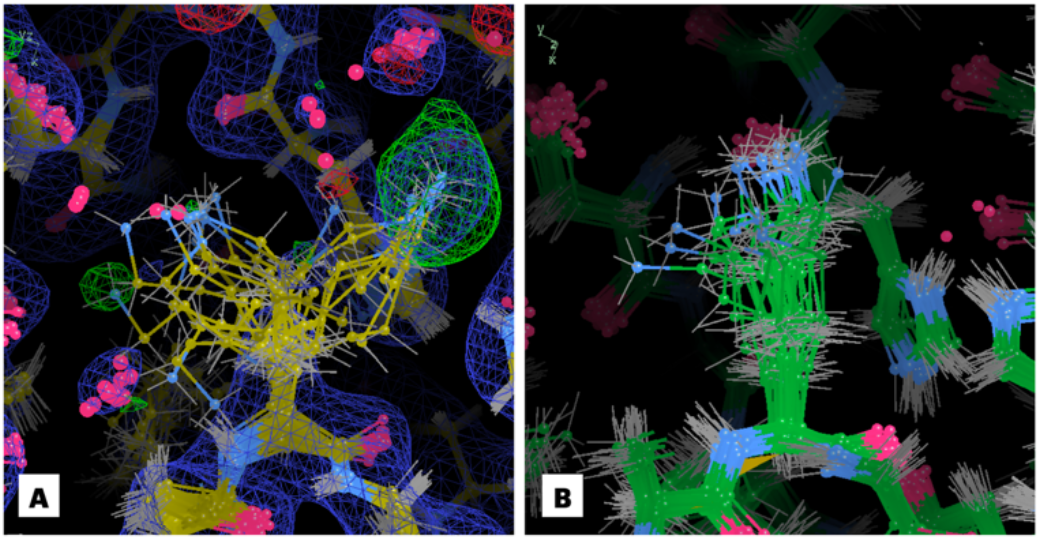
MD snapshot guides remodeling of Lys217. **A**: Coordinates, 2F_o_-F_c_ (blue, 1 sigma isosurface) and F_o_-F_c_ (positive in green, negative in red, 3 sigma isosurface) from ensemble refinement model *E*: the Lys217 amino group is diverse and includes extensions into off-backbone density and positive difference density where the phosphate was placed in the *R_f_* structure (Fig. 3). **B**: Final frame MD snapshot: the side chain conformations are more tightly clustered, extending straight from the backbone.

The water building step yielded a different side chain conformation for Lys217. The 1-sigma MD density envelope for the protein extends perpendicular to the backbone, and the conformation of Lys217 in *M*_all_ lies within this envelope (Fig. 3B). The side chain atoms in the *R_i_* structure are consistent with this conformation (Fig. 3C). The snapshot from the final frame of MD is consistent with the extended side chain conformation, and there is less variation among the structures from the snapshot than there is in *E* (Fig. 4B). None of the conformations in the snapshot are close to that in *S*, and none are close to those that fall within the extra density occupied by the amino groups in *E*.

The protein remodeling step provided a path forward for modeling the extra density near Lys217. Envelopes from both the isolated solvent and chloride MD densities overlapped the extra density, at first suggesting that it might include contributions from both (Fig. 3B). However, adding a water or chloride left a significant amount of positive difference density in the region, indicating that more electrons were required (Fig. 3C). This finding, along with the shape of the difference density, suggested the possibility that it might correspond to a free phosphate. Although free phosphate was not included in the simulation, phosphate was in the buffer, and at the experimental pH (< 6.5), these would be negatively charged, like the chloride. We therefore placed a phosphate into the difference density. In the resulting structure *R_f_*, the phosphate density appears reasonable, and the difference density is reduced to below the 3-sigma level (Fig. 3D).

Protein remodeling also yielded a multi-conformer model of Asp166. There is substantial difference density in the active site near MG1 in *S*, near Asp166 (Fig. 5C). There is also some positive difference density on either side of the side chain of Asp166: next to one of the oxygens on the α-carboxylic acid, and next to the Cβ-Cɣ bond (Fig. 5A). In *E*, Asp166 shows limited structural heterogeneity, with most of the side chains positioned as in *S* (Fig. 6A). In the final-frame MD snapshot Asp166 is similar to *S* in about half the structures; in the other half, however, it is shifted toward MG1 (Fig. 6B). The MD density also suggested a coordinated water adjacent to the oxygen on the α-carboxylic acid, and other waters next to MG1 (Fig. 5B).

**Figure 5:**
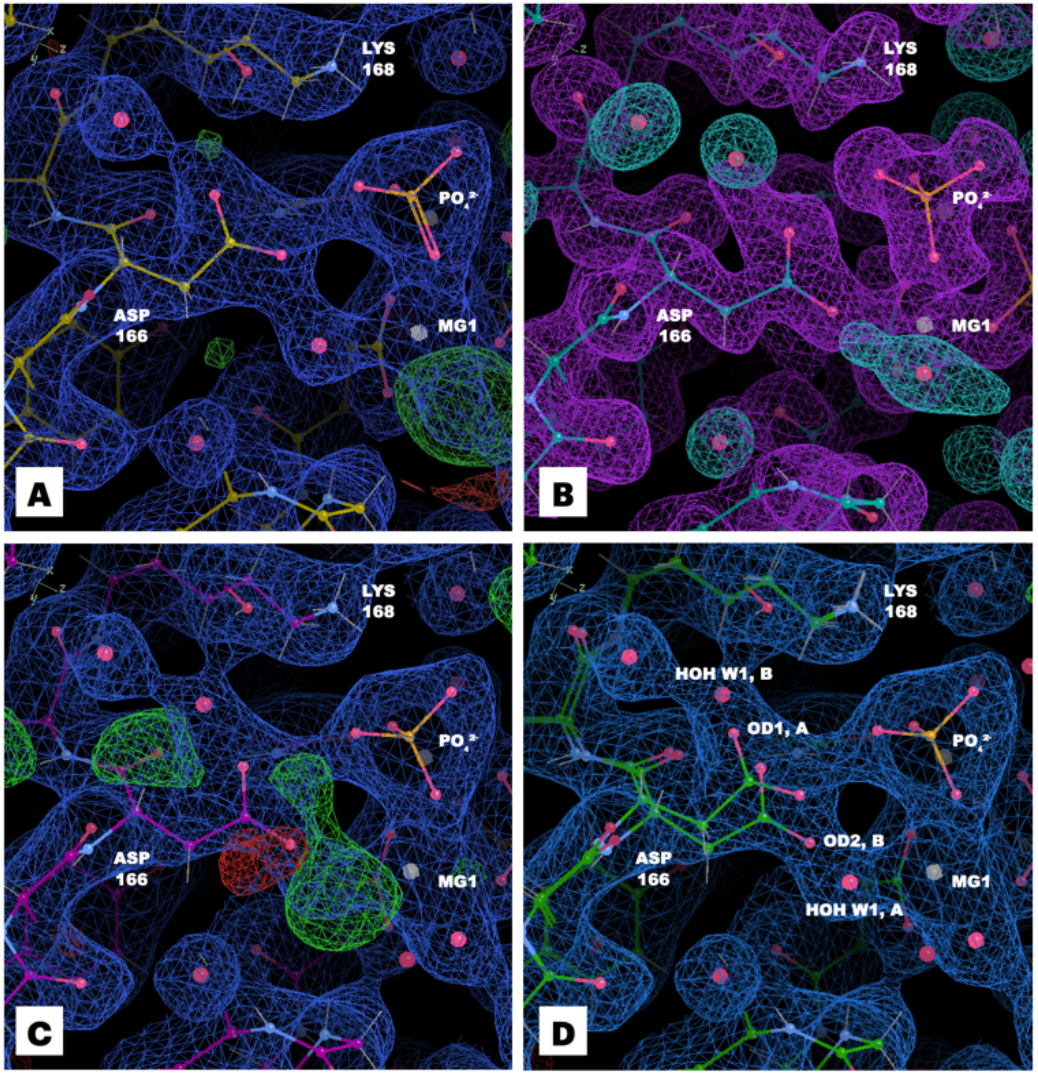
MD remodeling improves interpretation of density in the neighborhood of aspartic acid 166. **A:** coordinates, 2F_o_-F_c_ (blue, 1 sigma isosurface) and F_o_-F_c_ (positive in green and negative in red, 3 sigma isosurface) from model *S*. **B:** coordinates from the *M*_all_ model, with MD protein and cofactor density (purple, 1 sigma isosurface), solvent density (blue, 3 sigma isosurface), and chloride density (yellow, 10 sigma isosurface) from the 90-100ns segment of the 200 kJ mol^-1^ nm^-2^ simulation. **C:** coordinates, 2F_o_-F_c_ (blue, 1 sigma isosurface) and F_o_-F_c_ (positive in green and negative in red, 3 sigma isosurface) from model R_i_. **D:** coordinates, 2F_o_-F_c_ (blue, 1 sigma isosurface) and F_o_-F_c_ (positive in green and negative in red, 3 sigma isosurface) from model *R_f_*; water associated with Asp166 (labeled HOH W1, A/B) is modeled as a multi-conformer residue, with the A conformer adjacent to the magnesium, and the B conformer adjacent to the A conformer of the side.

**Figure 6:**
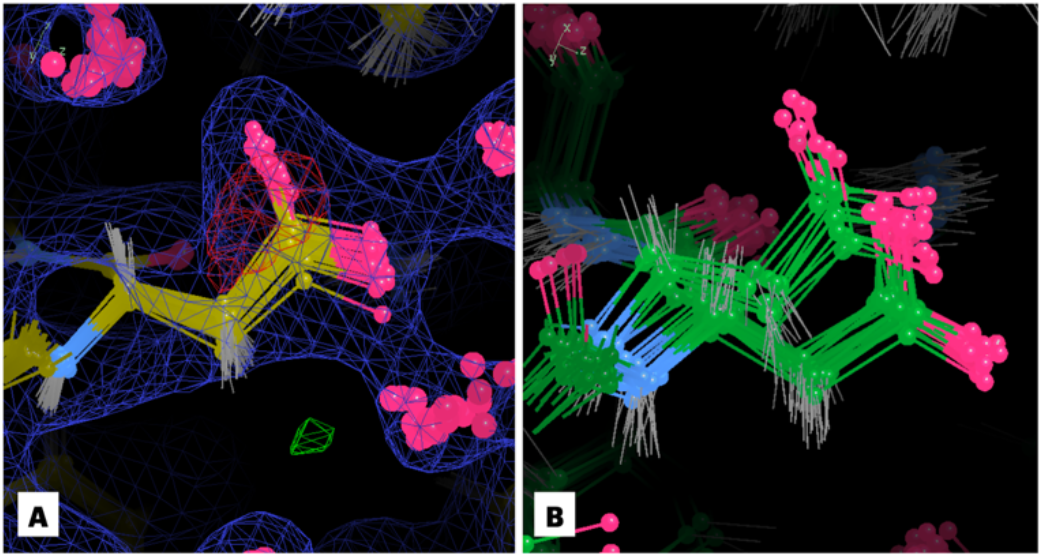
MD snapshot suggests multi-conformer model of Aspartic Acid 166. **A**: coordinates from ensemble refinement against experimental data, with 2F_o_-F_c_ (blue, 1 sigma isosurface) and F_o_-F_c_ (positive in green, negative in red, 3 sigma isosurface) from ensemble refinement. **B**: coordinates from the reverse-propagated final frame of the 200 kJ mol^-1^ nm^-2^ crystalline MD simulation. The MD ensemble exhibits significantly more structural heterogeneity than the ensemble from refinement, with about half of the side chains in the A conformation and half in the B conformation.

Based on the MD snapshot, we revised the model using alternate conformations for residues 163-169 (three residues to either side Asp166, allowing the backbone and side chain to shift to place the second conformation). At first it was not obvious how to model the water, because the MD water density appeared to clash with the multi-conformer model (Fig. 5C). To resolve this issue, a single water molecule was modeled with alternate conformations on either side of Asp166. In the resulting structure (Fig. 5D), the A position of HOH W1 is close to the B position of Asp166 OD2, and the B position of HOH W1 is close to the A position of Asp166 OD1; however, the clashes are avoided if the A and B conformations of Asp166 co-occur with their corresponding waters.

The revised model was refined against the crystallographic data, yielding *R_f_*. In *R_f_*, the density near Asp166 is substantially improved compared to *R_i_*, with nearly all 3-sigma difference density eliminated (compare Figs. 5C and 5D). In addition, the occupancies of the alternate conformations for residues 163-169 are consistent with the water occupancies: the protein and water A conformation occupancies are 0.56 and 0.51, respectively; and the protein and water B conformation occupancies are 0.44 and 0.49, respectively.

Like Asp166, protein remodeling produced a multi-conformer model of Lys213. The initial MD-revised model (*R_i_*) showed negative difference density close to the backbone oxygen of Lys213 and a large spot of positive difference density on the other side of the backbone (Fig. 7A). There was also a close contact (1.97 Å) between the backbone oxygen of Lys213 and a water oxygen (O/HOH/56/S) nearby. These features initially suggested the possibility of a peptide flip; however, substantial difference density remained after performing the flip (not shown). An inspection of the final-frame MD snapshot revealed a clear multi-conformer state for Lys213, with about half the structures occupying each of the peptide plane configurations (Fig. 7D). These states could not be identified clearly in *E*, which is much more disordered (Fig. 7C). The model was revised to include a multi-conformer state for residues 212 to 214 (*R_f_*) with the B conformation taken from one of the MD snapshot structures, reducing the difference density (Fig. 7B). The A conformation has occupancy 0.57, the B conformation has occupancy 0.43, and the water oxygen nearby is at occupancy 0.9 (compared to 1.0 in *R_i_*). The B conformation corresponds to a peptide-plane flip, also termed tt+ (34).

**Figure 7:**
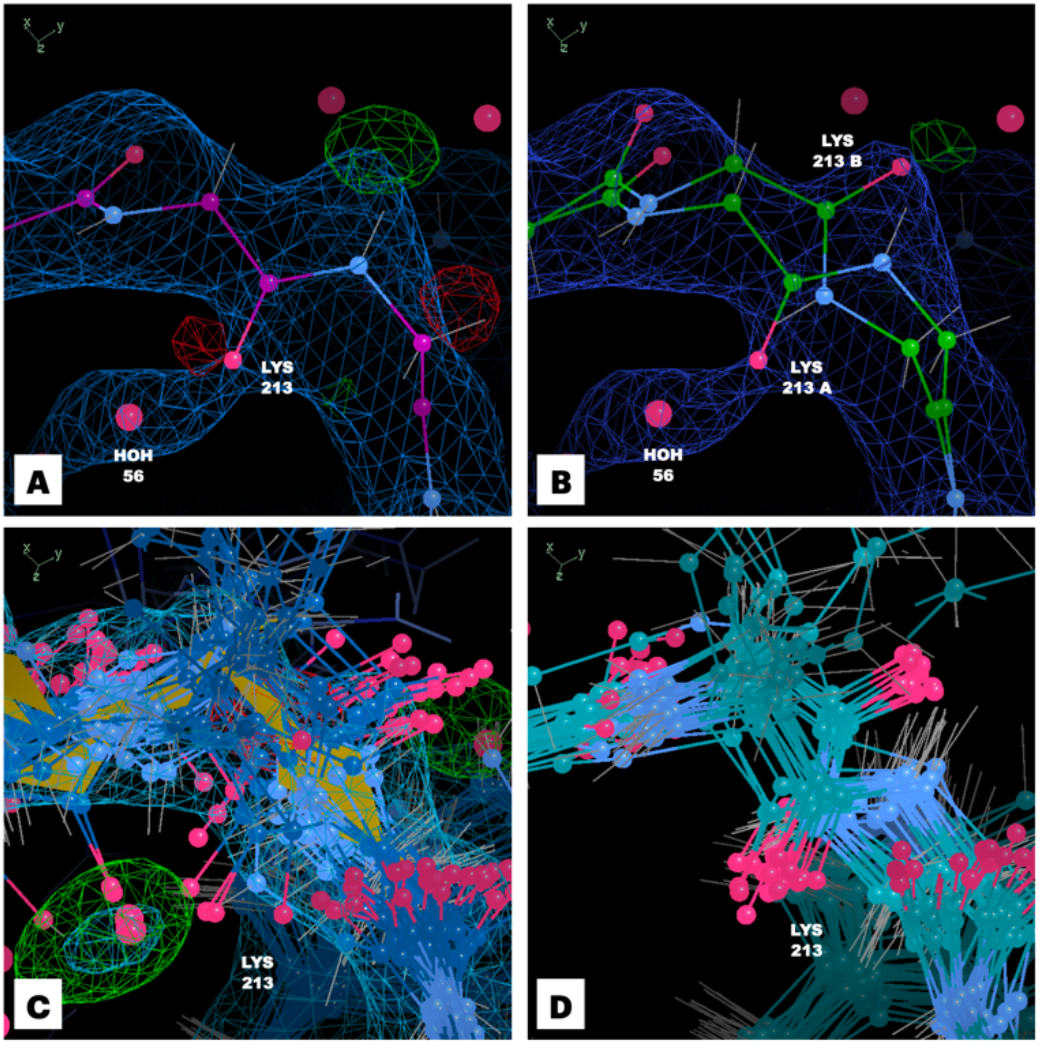
Density analysis and MD snapshot ensemble suggests multi-conformer state for Lys213. **A.** Initial MD-revised model (*R_i_*) coordinates (magenta), 2F_o_-F_c_ density (blue, 1 sigma isosurface) and F_o_-F_c_ (positive in green, negative in red, 3 sigma isosurface). **B.** Final MD-revised model (*R_f_*) coordinates (green), 2F_o_-F_c_ density (blue, 1 sigma isosurface) and F_o_-F_c_ (positive in green, negative in red, 3 sigma isosurface) — A conformation at 57% occupancy, B conformation at 43% occupancy, water oxygen O/HOH/35/S at 90% occupancy. **C.** Coordinates from ensemble refinement model (*E*) (blue), 2F_o_-F_c_ density (blue, 1 sigma isosurface) and F_o_-F_c_ (positive in green, negative in red, 3 sigma isosurface). **D.** Coordinates from ensemble snapshot from MD simulation (turquoise) — ensemble snapshot suggests a clear multi-conformer state defined by a peptide flip.

## Discussion

Computing crystallographic densities is a key element of the MD-MX procedure, as it enables the simulations to be compared directly to diffraction data. Such density comparisons enabled us to identify two residues whose conformations are especially sensitive to the protonation state: His62 and His294. Similar techniques may be used to falsify protonation states in other systems, and this approach also may be extended to investigate the impact of protonation on local ordered water structure and interactions with neighboring residues. Structural ensembles and calculated densities or structure factors from MD simulations may also prove useful for optimizing MD force fields using crystallographic data.

In addition to providing direct comparisons to the data, densities also provide a general approach to visualize structural ensembles that can complement other types of visualizations. For example, over the course of our study, we found that two sets of ensemble coordinates may appear different, while their average densities are the same. The converse may also occur: two sets of ensemble coordinates may be so disordered that it is hard to tell if there are any major structural differences, whereas their average densities may be obviously different. The latter case is particularly relevant for the study of water structure. Indeed, a recent solution-state MD study of protein water rehydration used density computations to overcome difficulties in using atomic coordinates directly to assess whether binding sites of interest were occupied by waters (35). Density comparisons like those performed here therefore should prove useful in MD applications beyond crystallography.

The MD-MX solvent model includes many ordered waters that were not present in the initial crystal structure. The modifications to the water network appear to be plausible in local regions of the protein that were examined in detail. For example, we removed waters from the active site and calculated a Polder map (an OMIT map without bulk-solvent correction) for the region; this density was extended and lacked strong peaks, which might help to explain why water picking did not place ordered waters in this region in the initial structure *S*. Nearly all the MD waters around the active site were supported by Polder density. Many of them were also near waters in a different structure obtained using neutron diffraction data (29). At present, we cannot say how many of the MD waters outside of the regions we examined carefully should be treated with high confidence. In some cases, there are waters with relatively weak support from the density compared to what one is used to seeing in crystallography. There are even some isolated cases where the waters are associated with negative difference density after refinement against experimental data; nevertheless, we chose to retain these waters, because our aim was to assess a water model entirely determined by the MD simulations, and to determine what we can learn about the crystal structure from such a model.

The MD-MX procedure helped to mitigate some of the pitfalls of protein crystallography that result from the inability to distinguish between protein and solvent density. Both single-structure and ensemble refinement may model side chains into total density which would otherwise be ambiguous as to the side chain’s conformation and/or identity (protein, solvent, or other molecules). The water building and protein remodeling methods provided information to resolve the ambiguities. The key information used in protein remodeling, beyond what is used in water building, comes from MD snapshots (Figures 4 and 6). In particular, the MD snapshots allowed us to identify three places where the crystal structure could be improved in ways not suggested by ensemble refinement: Lys217, Asp166, and Lys213.

Lys217 provides a clear example of how MD snapshots enable the disambiguation of density. In the initial crystal structure, *S*, there is a large amount of both positive and negative difference density in the region, and it is unclear how to remodel the side chain (Figure 3A). The ensemble structure, *E*, at first appears satisfactory, as much of the difference density in the region is reduced. In several of the structures, the side chain amino groups for Lys217 were placed into a spot of strong off-backbone density, and the remaining structures exhibit significant structural heterogeneity. However, there is still substantial positive difference density, suggesting that the model might still be improved (Figure 4B).

The protein remodeling method provided a clear path to improving both the side chain and the off-backbone density. Even though the simulation is restrained to the diverse set of structures in model *E*, the conformation of Lys217 in the MD snapshot was relatively homogeneous (Fig. 4B). This means that the force field was strong enough to overcome the restraints to the ensemble positions. After correcting the conformation of Lys217, difference density remained, and the MD simulation again guided the approach. Density comparison revealed that both MD water and chloride ion density overlapped the difference density. Although water and chloride atoms lacked sufficient electrons, when a phosphate was built in, this difference density was reduced to below 3-sigma. This free phosphate is not observed in the joint X-ray and neutron diffraction dataset, where there is a water molecule in the same position instead. However, for that experiment, phosphate was not present in the crystallization buffer, Sr^2+^ was used rather than Mg^2+^ to co-crystallize, and, although ADP is present, the gamma phosphate of ATP is transferred to a serine on the substrate peptide (29). The advantages seen in this example might be generalizable to other MX studies: charged ions in the solvent, which are often required for neutralization or modeling of buffer salts in MD simulations, may serve as proxies for other charged molecules in the experimental buffer that are not included in the explicit solvent of the simulation.

In addition to guiding the remodeling of a single conformation of a side chain, as for Lys217, the cases of Asp166 and Lys213 show that the MD-MX procedure can help develop multi-conformer models. Although in retrospect the difference density from single structure refinement contained clues about the potential for a peptide-plane flip for Lys213, the possibility was only suggested after producing an alternative water model from the MD simulations: when the resulting model was subjected to comprehensive validation, a water oxygen was flagged as being too close to the backbone oxygen of Lys213. It was this close contact that drew our attention to the difference density in the region. Because a simple peptide flip did not address the issue, we attempted to use the ensemble refinement for guidance about how to revise the structure; however, the conformations of Lys213 in *E* were too disordered to suggest a path forward. In contrast, the MD snapshot exhibited a clear two state conformational distribution for the backbone that we were able to use to define an alternative conformation (Fig. 7). Similarly, though both the single structure and ensemble refined models for Asp166 have difference density in the region of the side chain, neither suggests a path forward — again, the MD snapshot clearly suggested a specific multi-conformer model (Fig. 6). These examples show that the MD-MX procedure is useful not only as an analysis tool, but also can be used to develop multi-conformer models of protein structure, including peptide flips (34, 36).

A key step in the MD-MX procedure is the refinement of structure *S* into the MD structure factors calculated from just the protein atoms (“protein-first” refinement). To assess the importance of this step, we altered the procedure by skipping it and instead directly refining *S* against the MD structure factors calculated from the entire system. The resulting model differed in three key ways from the one resulting from protein-first refinement. (1) There are isolated cases where side chains are built into water density (for example, Lys28 and Lys295). As refinement against experimental data is always performed using the total structure factors, such errors might not be uncommon in published crystal structures. (2) The R-factors obtained after initial refinement of this structure against the experimental data (R_*i*_) are higher (R_work_ of 0.1382 rather than 0.1328 and R_free_ of 0.1795 rather than 0.1777), indicating that protein-first refinement improved the agreement with the data. (3) The model contained about 50 fewer waters. Inspection of the missing waters revealed that they are primarily in the second hydration layer and beyond (not shown). The overall shape of the water density envelope appeared to be similar with or without protein-first refinement; the differences nevertheless influenced the water picking, perhaps during the filtering step.

Our study yielded substantial insights into PKA structure and function. The PKA-C crystals studied here were prepared in a relatively uncommon state: with both crystal preparation and diffraction performed at room temperature. The room temperature preparation allowed us to capture an intermediate catalytic state with ATP hydrolyzed to ADP, with the free phosphate and two magnesium ions bound in the active site. The active site has previously been observed with ATP hydrolyzed in a mutant (37). A neutron diffraction structure (29) captured the active site in a similar state, but with the phosphate transferred to a residue on a bound peptide. The present structure is unique in that the free phosphate is present in the active site. In this structure we have thus captured the slow ATPase activity of the C-subunit which leads to a reduced affinity for the inhibitor peptide (38).

The sensitivity of the conformation of His294 to the protonation state was notable: when doubly protonated, the side chain of His294 entered an entirely different rotameric state, making room for two ordered waters to enter the space occupied by the side chain when singly protonated. This sensitivity might be biologically significant: previous studies have shown that this region, and this residue in particular, plays a role in the binding and release of substrate, with mutations of residues in this region modulating the affinity of binding (33, 39). His294 might become doubly protonated when the molecule is brought close to charged groups; for example, the regulatory subunit PKA-RII binds to the membrane, and when bound, would position PKA-C such that this region would be near phospholipid head groups (40, 41).

Previous studies have shown that both Lys213 and Lys217 are crucial for binding to the regulatory subunit – with charge-to-Ala mutations for both residues having a significant effect on binding affinity to RIɑ (42). The extended conformation of Lys217 and the addition of the free phosphate in the region may have implications for binding to the regulatory subunit. At present, we are not aware of any biological role this free phosphate may play; it would be interesting to determine whether it might have a role in regulating activity. The region surrounding Lys213 is a basic surface (PRS2) involved in the binding to PKA-R. In 2QCS (42), Lys213 makes a crucial link with Thr237 and Arg241 of RIɑ (Figure 8), which together form an allosteric hotspot that coordinates communications between two cyclic nucleotide binding domains (CNB-A and CNB-B). The multi-conformer state for Lys213 suggests that conformational selection might be important in binding of PKA-C to PKA-R: while previous models of this region have the residue in a single conformation for either the bound or unbound states, our structure suggests that the side chain naturally samples two conformations at room temperature (Fig. 7).

**Figure 8:**
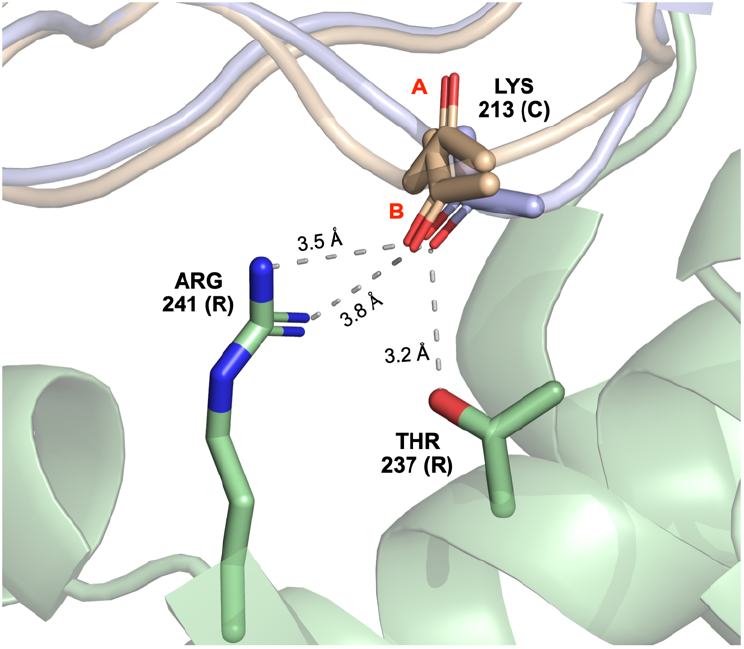
Alternative conformation of Lys213 is consistent with backbone pose for binding to regulatory subunit. Final MD-revised structure (*R_f_*, light brown) models Lys213 of the catalytic subunit (PKA-C) with two conformers defined by a peptide flip (side chain not shown). The B conformer of Lys213 is consistent with the backbone oxygen orientation of PKA-C bound to the regulatory subunit, RIɑ (pdbid 2qcs shown with catalytic subunit shown in light blue and regulatory subunit shown in light green). The backbone oxygen of Lys213 (C) is close to Thr237 and Arg241(R).

We are unaware of a multi-conformer model for Asp166 having been suggested before in the literature. It is possible that the presence of this state is connected to the hydrolyzed ATP and the free phosphate, and the result of our room temperature crystal growth along with data collection at 12°C. This model of Asp166 suggests a mechanism of activity involving the water network adjacent to MG1 (Fig. 9). When Asp166 is in the A conformation, the distance between MG1 and the water oxygen HOH W1A is 2.12 Å. In the B conformation, this water is displaced, and the MG1 is coordinated with OD2 of Asp166 with a longer distance of 3.00 Å. The change in coordination distance for MG1 going from the water oxygen to the oxygen of Asp166 suggests the possibility that the placement of MG1 might be less energetically favorable when Asp166 is in the B conformation, compared to the A conformation. Previous studies have shown that simultaneous release of both magnesium ions and ADP from the active site is unfeasible, and instead, one magnesium (MG1) must first exit before ADP and MG2 can exit, post-catalysis (43, 44). The multi-conformer states for both Asp166 and associated water(s) thus might correspond to a progression of the phosphotransfer reaction and suggest a potential mechanism for this postcatalysis release.

**Figure 9:**
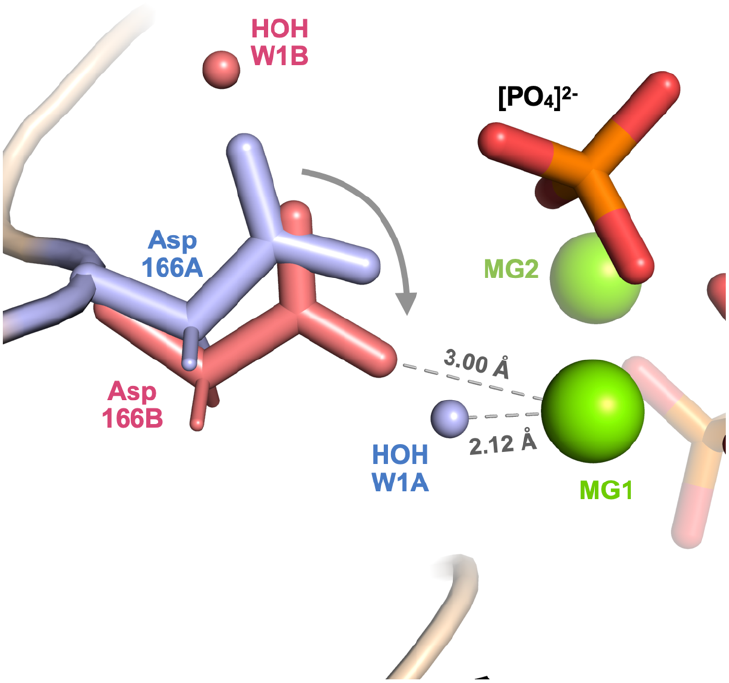
Implications of the Full-MD Revised model, *R_f_*, for structural changes associated with catalysis. In the A conformation (blue), Asp166A is shifted away from MG1, and water oxygen HOH W1A is coordinated with MG1. In the B conformation (red), Asp166B is shifted down toward MG1, and makes room for water oxygen HOH W1 to occupy a different site above the side chain (where it would have clashed with conformation A for Asp166). The distance between oxygen OD2 on Asp166B is slightly larger (3.00 Å) than the distance between water oxygen HOH W1A (2.12 Å); potentially limiting coordination with MG2 and encouraging its escape from the active site post catalysis. The multiple conformations suggest the possibility of a concerted motion (arrow) for Asp166 associated with the progression of catalysis.

The insights gained into PKA suggest that the MD-MX procedure may be beneficial in other crystallographic studies, producing information that cannot be obtained from other methods: (1) The MD-MX procedure yields an MD-derived solvent model that can be compared to the one generated by water picking in crystallographic refinement; (2) The MD solvent model and density analysis can help to guide manual adjustments when the difference density does not provide a clear picture of how to improve the structural model; (3) It yields picture of conformational variability that is more independent of the crystallographic data and is more integrated with the solvent model, in comparison with other techniques. For example, *qFit* (1, 3) and *phenix.ensemble_refinement* (6) generate pictures of anharmonic motions that go beyond what is possible using B-factors, and also can yield improved R factors. However, *qFit* does not attempt to model solvent density, and *phenix.ensemble_refinement* does not model all waters explicitly in the MD simulation step. Similarly, other non-MD based ensemble refinement approaches (4, 5, 45) do not consider the solvent in exploring multiple conformations. These limitations can lead to placement of side chains into non-protein density, as we saw in the case of Lys217 (Fig. 4). In contrast, the MD-MX procedure can provide information to distinguish protein and solvent density. The MD-MX procedure also provided a clear model of multi-conformer states for residues for Lys213 and Asp166. In contrast, the information from the ensemble refinement model was unclear for these cases: in the case of Lys213, the conformations are highly disordered, without a clear two-state distribution; and in the case of Asp166, there is limited conformational variability in the ensemble refinement model, and there is substantial difference density. It is possible that differences in the MD simulations between the methods (explicit solvent, crystalline simulation, the force field, etc.) contribute to the difference between the MD-MX method and ensemble refinement. Even when other techniques may suggest similar remodeling ideas, the MD can provide independent information to increase or decrease confidence in these ideas.

To apply the MD-MX procedure more broadly, it will be useful to develop it into an automated workflow. Such a workflow could then be folded into the crystallographic refinement pipeline in such a way as to iteratively alter and improve the ordered water and protein structure, similar to, e.g., real space refinement or water picking and filtering. Certain aspects of the procedure lend themselves quite naturally to automation, such as the preparation and performing of the MD simulation and the calculation of densities. Other parts of the procedure will be more challenging to automate, such as multi-conformer modeling when there are potential water clashes.

## Materials and Methods

### Crystallographic Data Collection and Refinement Details

The C-subunit of PKA was recombinantly expressed in *E. coli* and purified as previously described (46). An additional MONO-S cation exchange step was used to separate phosphorylation states, with the peak corresponding to three phosphorylations taken for crystallization. C-subunit was concentrated to 8 mg/ml in 0.02 M KH_2_PO_4_ pH 6.5, 0.15 M KCl, and 0.001 M DTT and a ternary complex with inhibitor and ATP was formed with 1:20:10:10 molar ratio of C-subunit:Mg:ATP:IP20. The complex was crystallized by hanging drop vapor diffusion at 20°C under conditions as previously described (47) in 20% PEG 4000, 0.05 M MES pH 5.2, 0.05 M MgCl_2_, and 0.005 M DTT.

Room temperature (12°C) X-ray diffraction data were collected at beamline 8.3.1 at the Advanced Light Source (ALS). The crystal was mounted on a MiTeGen^™^ loop and sealed in a MicroRT capillary. Two sweeps of 180° were collected between translations along the crystal. The sweeps were processed independently with a resolution cutoff of 2.4 Å using DIALS/xia2 in ccp4i2 (48, 49). BLEND in ccp4i (50) was then used to remove frames with radiation damage, merge, and scale the dataset (merging statistics in Table S1). Molecular replacement was performed using *phaser* (51) with 3FJQ (52) as the search model.

Refinement and model building was performed in *phenix* (53) and *coot* (54). The input structure for ensemble refinement was prepared using an initial single-structure refinement model with full occupancies for main chain and inhibitor peptide atoms, refined occupancies for the phosphate, ADP, and magnesium ions in the active site, and refined sites and B-factors for all atoms, as well as ordered waters placed by water picking, producing a model (*S*) with R_work_ = 0.1557, R_free_ = 0.1838. The final model had 148 waters. Due to a lack of supporting density, this model had 14 residues missing in the flexible N-terminus, and atoms missing in residues Gln176, Glu248, and Lys254. The waters around the active site of this model were removed, and an OMIT map was created, eliminating bulk solvent correction, to create a “Polder” map (55), which represents the density due to the water atoms in the region that were not included in the model.

An ensemble structural model, *E*, was refined with *phenix.ensemble_refinement* (6) for use in the crystalline MD. The number of ensemble members was set to 32 (one for each protein in the supercell system) by setting *ensemble_reduction* to “False”, and restricting the *number_of_acquisition_periods* and *pdbs_per_block* to 8 and 4, respectively (8 x 4 = 32).

### MD System Preparation

Previous crystalline MD simulations have used a single structure refinement model for the initial coordinates for all proteins in the supercell. Here, we used a modified *E* model and seeded each protein in the supercell with a unique conformation from the ensemble. Because model *E* had 14 residues missing on the flexible N-terminus as well as residues 23-24 of the PKI peptide not modeled due to poor density, to produce the model which was used to seed the crystalline MD supercell, these residues were modeled in from 1CMK (56) into the 32 structure ensemble and the added residues had B-factors set to 120 and were refined in Phenix using rigid body and real-space refinement followed by geometry minimization. No steric clashes were observed for the propagated system, and no issues were encountered in energy minimization, solvation, or equilibration.

After removing crystallographic waters, each of the structures from the ensemble was propagated to a different location in a 2×2×2 unit-cell supercell according to the symmetry information provided by the CRYST1 record (using two custom Python modules, *pdbio.py* and *propagate.py*, available at*https://github.com/lanl/lunus/scripts*). The full system was parameterized in AMBER’s *tleap* (from the *ambertools* package version 20.14) (57) using the AMBER14SB (58) force field for the protein and magnesium ions and the *phosaa10* parameter set (59) for the phosphorylated serine (SEP) and tyrosine (TPO) residues. The active-site-bound ADP was parametrized with the *frcmod* and *prep* files found in the Bryce Lab AMBER parameter database for cofactors (60). The phosphate was assumed to be doubly protonated ([H_2_PO_4_]^-^) and was subjected to *mp2/aug-cc-pvdz* QM geometry optimization to determine the geometry of the doubly-protonated state, before being parametrized in *tleap* with the *gaff2* force field. The phosphate was placed identically into each ensemble structure with least-squares distance to the crystal structure phosphate molecule minimized, and hydrogens oriented away from the active site magnesium ions.

The protein model was prepared in two alternative protonation states. In one model, every histidine was doubly protonated (HIP); in the other model, the protonation/tautomeric states were assigned based on data from neutron diffraction: histidines 87, 158, and 260 remained doubly protonated (HIP), histidines 62, 131, 142 and 294 were protonated on the epsilon nitrogen (HIE) and histidine 68 was protonated on the delta nitrogen (HID) (29). The system was initially solvated (using GROMACS’ *solvate* (61)), neutralized with chloride ions (using GROMACS’ *genion*), and solvent atoms were replaced with magnesium and additional chloride ions sufficient to mimic the crystallization buffer MgCl2 concentration of 0.05 M.

### Solvation and Equilibration

Standard MD equilibration takes place in the NPT ensemble, in which the box side lengths are allowed to fluctuate to maintain constant pressure. However, to minimize errors in computing mean structure factors, it is important for us to perform simulations with the periodic box side lengths fixed — i.e. in the NVT ensemble (21). Upon initial solvation and equilibration, the system was dramatically under-pressurized (about −1000 bar). To bring it up to atmospheric pressure, the system was subjected to iterative rounds of additional solvation, minimization (using the steepest-descent algorithm), and NVT equilibration (500 ps total, with restraints on all heavy atoms with restraint constant equal to 200 kJ mol^-1^ nm^-2^), until the average measured pressure, plus or minus standard error, was in the range of −100 to 100 bar. The system was then simulated for 10 ns of restrained “preliminary production” at the same restraint strength, to ensure that the system was fully equilibrated, before the system was simulated for another 100 ns of “production” trajectory.

### Production Simulation

Both the HIP-protonation model and the model with protonation states based on neutron diffraction were simulated with position restraints on heavy atoms using a spring constant of 200 kJ mol^-1^ nm^-2^. We also performed an additional unrestrained simulation. In all simulations, the time step was 2 fs (with LINCS constraints on hydrogen bonds) with coordinate output every 2 ps. During the early portion of the trajectories, the RMSD of atom positions between the MD protein structure and the crystal structure decreased steadily, under the influence of the restraints. In the restrained simulations, a plateau was reached at about 60 ns, and the 90-100 ns portion of the production trajectory was used for density analysis. The trajectory was down-sampled to 1 frame every 10ps and processed to keep molecules whole and account for periodic boundary corrections (using GROMACS’ *trjconv* method with flags *–pbc mol* and *-pbc nojump*). Additional details on the simulation parameters are available in the *.mdp* files which are included in the Supplemental Materials.

### MD Density Analysis

Mean structure factor *.mtz* files were calculated from the final 10ns analysis trajectories using *xtraj.py* (23), which takes a GROMACS *.xtc* trajectory file and *.pdb* topology file (taken from the first frame of the simulation) as input. MD structure factor files were calculated separately for (1) the full system, (2) protein, (3) water, (4) magnesium, and (5) calcium atoms. The *fft* method from ccp4 (version 7.1(62)) was used to convert the structure factors to electron-density maps. The maps were placed on an absolute scale (*e*^-^ Å^-3^) using the unit-cell F_000_ and volume, which were calculated using *cctbx* methods (63).

The MD water density was analyzed with the *peakmax*, and *sftools* ccp4 methods to produce a set of peak positions with peak heights above a threshold of 1 electron per cubic angstrom, representing ordered water positions from the MD. The recall statistics for crystallographic waters were calculated by isolating the unique waters from the set produced by peak-finding (atom indices for symmetry-related copies have the same atom number), calculating the distance between crystallographic and MD-precited waters (modulo unit cell periodicity), and determining the fraction of crystallographic waters that have an MD-predicted water within a specified cutoff distance.

### MD Snapshots

Custom Python scripts (*pdbio.py* and *reverse_propagate.py*) were used to analyze the final frame of each simulation and compare the original crystallographic ensemble to the ensemble from the final frame of each MD simulation (available at https://github.com/lanl/lunus/scripts).

### MD-Informed Model Refinement

The Initial Single Structure Model refined against experimental data was stripped of its waters and refined (using *phenix.refine*) against the average structure factors from only the non-solvent components of the system with bulk-solvent corrections and water picking turned off. Real space refinement was activated, to allow side chains to find the conformations represented in the MD density. Next, bulk solvent corrections were turned on, and ordered water picking was activated for a single round of refinement against the full average structure factors from the MD to produce a new set of waters. The resulting structural model was then fed back into refinement against experimental reflection data without water picking, to produce a single structure model refined against the experimental data, with the protein structure and ordered water positions derived from the MD data (Initial MD-revised Model, *R_i_*, in Tables 1, S1). Side chains that were not initially modeled due to poor experimental density (Q176, E248 and K254) had missing atoms added with *coot*’s *Simple Mutate* method (54), and the sites and B-factors for the atoms in only these residues were refined for a single macrocycle. To incorporate the findings from MD (discussed above) a phosphate was added to this model in a location suggested by chloride ion density and a multi-conformer region was constructed for residues 163 to 169, and 212 to 214 with the A conformation pulled from the single structure refined against experimental data and the B conformation taken from the final frame MD reverse-propagated ensemble, and both conformations were set to occupancy 0.5. The A and B conformations for Asp166 both had associated waters in their respective single structure models, and each of those waters was given the same alternate conformation ID (A or B) as its associated side chain, and given occupancy 0.5. Additional waters were added around the active site for which both difference density and MD solvent density was present above a 3 sigma threshold. The resulting model was refined against the experimental reflection data for 3 macrocycles, yielding the final structure informed by the MD simulations (Final MD-revised Model, *R_f_*, in Tables 1, S1).

Crystallographic data and atomic coordinates for the final MD-revised structure *R_f_* have been deposited at RCSB with PDB ID 7UJX.

## Acknowledgments

We thank Jian Wu, Alexandr Kornev, and Jessica Bruystens for discussions. This work was supported by the University of California Laboratory Fees Research Program (No. LFR-17-476732). DCW and MEW are supported by the Exascale Computing Project (17-SC-20-SC), a collaborative effort of the DOE-SC and the DOE National Nuclear Security Administration. DCW is supported by the Center for Nonlinear Studies at Los Alamos National Laboratory. JSF is supported by NIH GM123159 and GM124149. PCA was supported by T32 CA009523/CA/NCI NIH HHS. SST is supported by NIH R35 GM130389. DLM is supported by NIH 1R01GM108889-01, 1R01GM124270-01A1 and R01GM132386.

---

**Supplementary Table S1:**
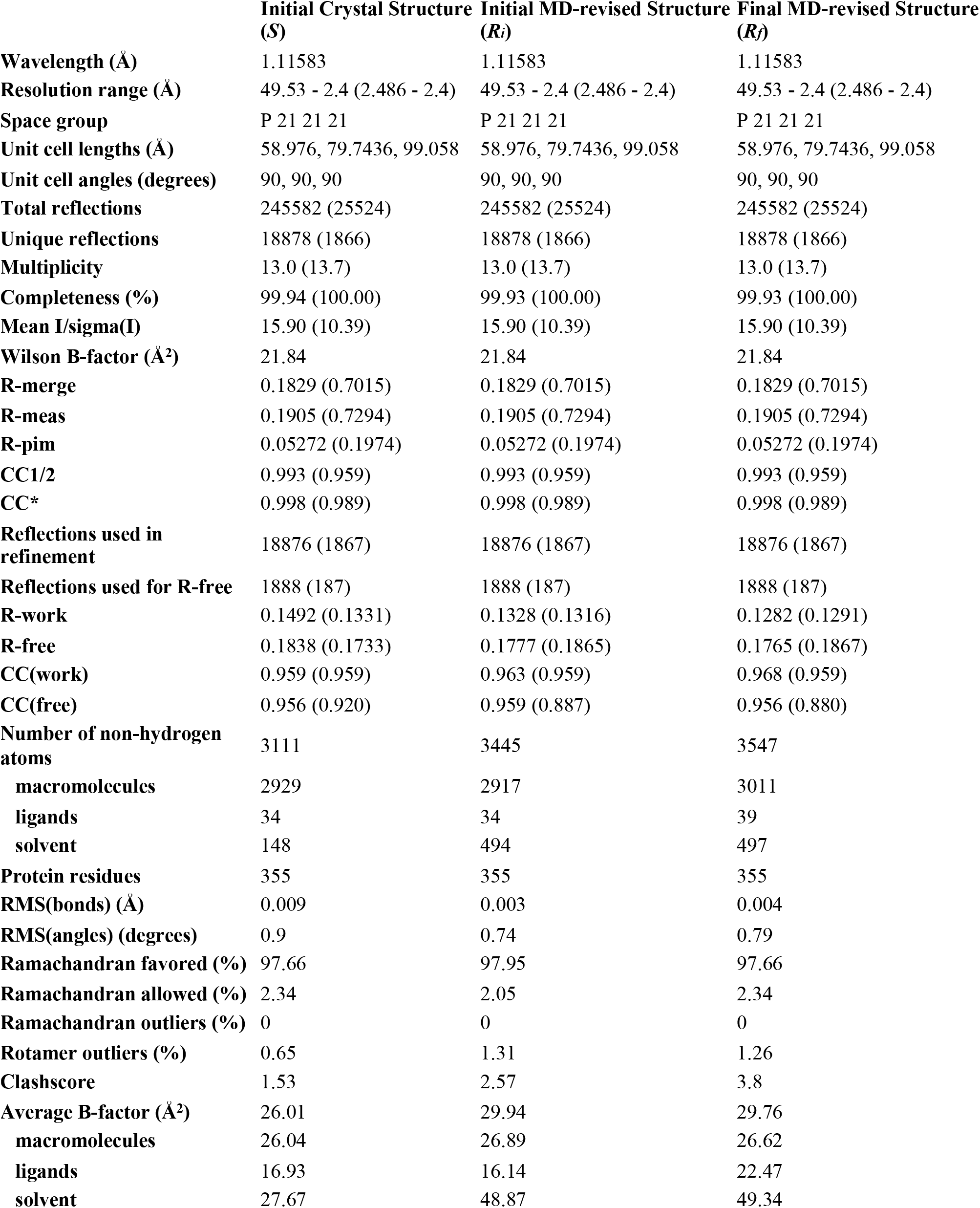
Crystallographic table one produced by *phenix.table_one*.

